# Cell-to-cell variability in inducible Caspase9-mediated cell death

**DOI:** 10.1101/2021.06.08.447627

**Authors:** Yuan Yuan, Huixia Ren, Yanjun Li, Shanshan Qin, Xiaojing Yang, Chao Tang

## Abstract

iCasp9 suicide gene has been widely used as a promising killing strategy in various cell therapies. However, different cells show significant heterogeneity in response to apoptosis inducer, posing challenges in clinical applications of killing strategy. The cause of the heterogeneity remains elusive so far. Here, by simultaneously monitoring the dynamics of iCasp9 dimerization, Caspase3 activation and cell fate in single cells, we found that the heterogeneity was mainly due to cell-to-cell variability in initial iCasp9 expression and XIAP/Caspase3 ratio. Moreover, multiple-round drugging cannot increase the killing efficiency. Instead, it will place selective pressure on protein levels, especially on the level of initial iCasp9, leading to drug resistance. We further show this resistance can be largely eliminated by combinatorial drugging with XIAP inhibitor at the end, but not at the beginning, of the multiple-round treatments. Our results unveil the source of cell fate heterogeneity and drug resistance in iCasp9-mediated cell death, which may enlighten better therapeutic strategies for optimized killing.

## INTRODUCTION

Inducible Caspase9 (iCasp9) is a cellular suicide gene that allows conditional cell elimination^1^. It comprises a human Caspase9 fused with an inducer-binding domain which could be dimerized by the Chemical Inducer of Dimerization (CID), AP20187 or AP1903^2,3^. iCasp9 is dimerized and activated by induced proximity of the inducer-binding domain, leading to activation of the apoptosis pathway^4^. Due to the efficient induction of cell death, iCasp9 has been used as one of the most promising killing strategies in cancer therapies^5–7^ and adoptive cell therapies^8–14^. However, cells, especially cancer cells, cannot be eliminated 100% even treated with an extremely high dose of drug. The incomplete killing of cells leads to drug resistance and hampers the further use of the iCasp9 suicide gene^15,16^. Consequently, one potential way to improve the performance of iCasp9 is through increasing drug efficacy. An understanding of the source of heterogeneous cell responses to the drug and how such heterogeneity contributes to resistance could lead to more effective treatment strategies.

Previous investigations into isogenic populations of tumor cells in response to apoptosis inducing drugs revealed that the heterogeneity could emerge from non-genetic mechanisms, often through stochastic fluctuations in protein expression^17–20^ or differences in dynamics^21,22^. These studies mainly focused on the upstream of the apoptosis pathway. However, in the iCasp9 system, CID triggers the very downstream apoptosis pathway, so the source of heterogeneous response can be different. As depicted in Fig. 1a, in response to CID, iCasp9 is first dimerized and activated through the proximity of DmrB protein. Then, active iCasp9 triggers the activation of the executor Caspase3, which cleaves thousands of substrates in cells^23^, resulting in irreversible cell death^24,25^. Opposing the pro-apoptotic iCasp9 and Caspase3, the anti-apoptotic protein XIAP inhibits and degrades pro-apoptotic caspases once they are activated^26^. Thus, cell death is regulated by interactions between pro-apoptotic and anti-apoptotic proteins.

**Fig. 1:**
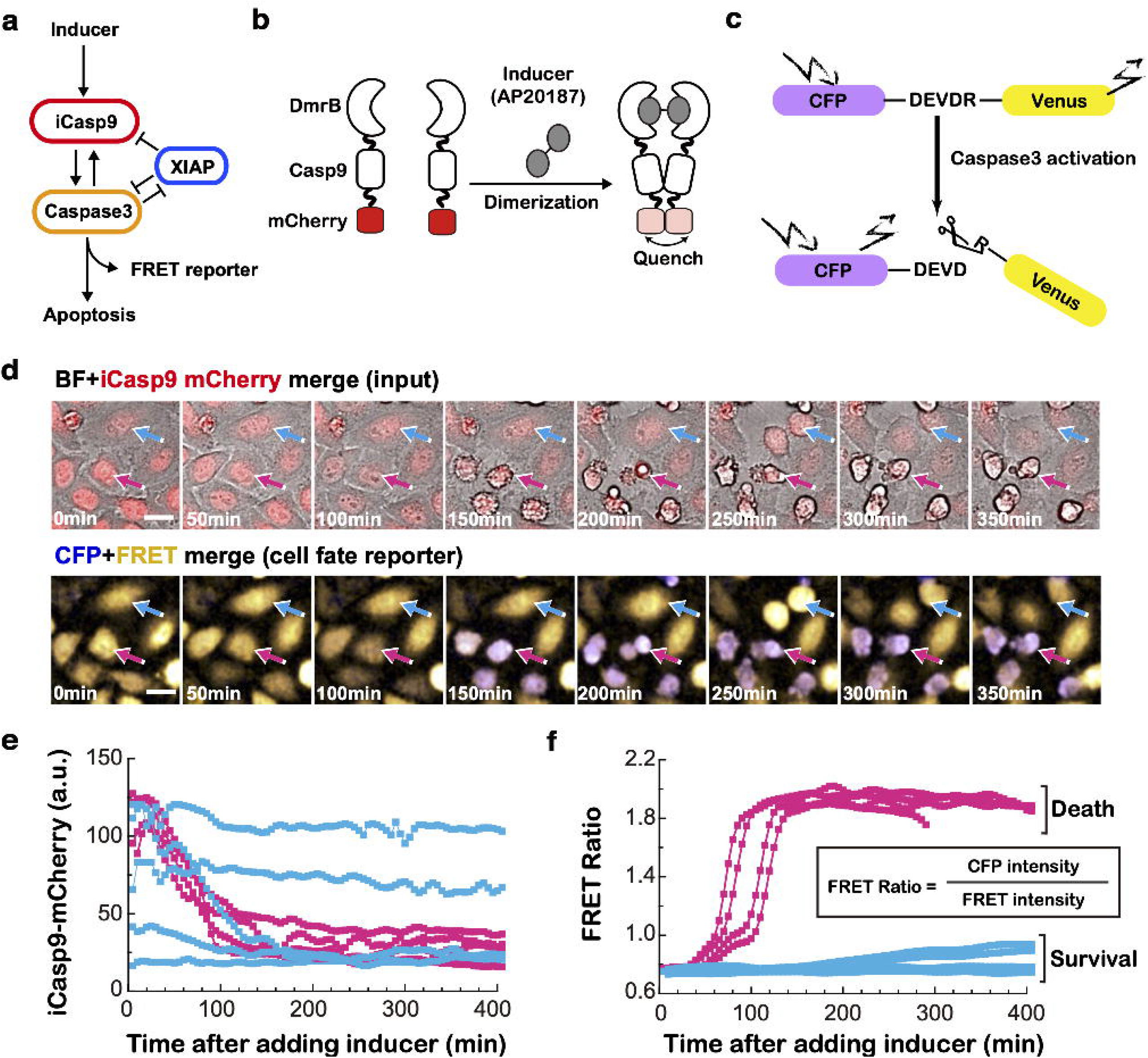
Construction of iCasp9 system to monitor iCasp9 dimerization and cell fate simultaneously in single cells. **a,** AP20187-induced apoptosis pathway in iCasp9 cells. **b,** Schematic diagram for induction of iCasp9 dimerization. White half-moon shape represents the DmrB domain which can be bound and dimerized by inducer (grey). White square represents full-length Caspase9, which is tagged with an mCherry fluorescence protein (red). Upon dimerization, the iCasp9-mCherry fluorescence signal will be greatly quenched. **c,** The FRET reporter indicating Caspase3 activity and cell fate. **d,** Microscopy images of merged BF and iCasp9 mCherry channel (upper panel) and merged CFP and FRET channel (lower panel) for iCasp9 cells treated with 0.25 nM AP20187. The blue and the pink arrow indicates a typical survival and dead cell, respectively. Images were taken under a 40× confocal fluorescence microscope. **e,** Single-cell profiles of iCasp9-mCherry fluorescence signals in dead cells (shown in pink) and survival cells (shown in blue) after addition of 0.25 nM inducer. **f,** Single-cell profiles of FRET Ratio signals in dead cells and survival cells after addition of 0.25 nM inducer.

In this work, we investigated potential causes for the heterogeneous cell response to inducer AP20187. We tagged iCasp9 with an mCherry fluorescence protein to track the dynamics of iCasp9 dimerization and incorporated a fluorescent substrate for Caspase3 to report Caspase3 activation and cell fate. We found that heterogeneous cell fates were originated from cell-to-cell variability in the initial iCasp9 expression and the ratio between XIAP and Caspase3 expression levels (XIAP/C3 ratio). Moreover, we also observed significant heterogenous behaviors within survival or dead cells, which were characterized by different dynamics of iCasp9 dimerization and Caspase3 activation within the survival cell population, and a broad distribution of death time within the dead cell population, respectively. Changing the inducer concentration not only changed the killing efficiency, but also dramatically altered the composition of survival cell types as well as the distribution of cell death timing. Additionally, multiple rounds of inducer application weakened the killing efficiency due to the accumulation of drug resistance originated from a selective pressure on low initial iCasp9 level. The killing efficiency on the resistant population could be greatly improved by the combinational use of the inducer with XIAP-targeted drug at the end of the multiple-round inducer treatment. Interestingly, the elevation of killing efficiency brought by the combinatorial drug use was much less effective if the drug combination was used from the beginning of the multiple-round treatment. Taken together, our results unveiled the source of cell heterogeneity in iCasp9-mediated cell death and demonstrated the “smart application” of combinational drugging for optimizing the killing efficiency, which may offer insights and new strategies for better clinical use of the iCasp9 system.

## RESULTS

### A system to monitor iCasp9 dimerization and cell fate simultaneously in single cells

To investigate what determines the heterogeneous responses of isogenic iCasp9 cells to the inducer, we first developed a system to simultaneously monitor iCasp9 dimerization and cell fate in single cells.

To monitor iCasp9 dimerization, we constructed an isogenic iCasp9 Hela cell line, in which iCasp9 was tagged with an mCherry fluorescence protein. As fluorescence signals can be greatly quenched upon dimerization due to the reduction of electronic energy level^27–29^, the dimerization of iCasp9 thus could be reflected by the decrease of mCherry fluorescence intensity (Fig. 1b). Indeed, we observed a significant fluorescence drop after adding the inducer (Supplementary Video 1), while no fluorescent intensity change was observed either in microtubule-mCherry cells with inducer addition (Supplementary Fig. 1a) or in iCasp9-mCherry cells without inducer addition (Supplementary Fig. 1b).

For the cell fate, since cells will all undergo apoptosis once Caspase3 is activated, we introduced a Fluorescence Resonance Energy Transfer (FRET)-based reporter, CFP-DEVDR-Venus, to monitor the Caspase3 activation (Fig. 1c)^30^. In this reporter, CFP and Venus are linked together by five amino acids ‘DEVDR’, which is the optimal substrate sequence for Caspase3. The ratio of CFP *versus* Venus fluorescence (FRET Ratio) will change upon cleavage of the linker ‘DEVDR’ (Fig. 1c and Supplementary Fig. 1c). The FRET Ratio of the reporter is highly correlated with the morphology change of apoptotic cell as we monitored cells under the microscope (Fig. 1d). While most unperturbed iCasp9 cells were alive with a low FRET Ratio (i.e. the FRET reporter within cells was intact without Caspase3 activation, Supplementary Fig. 1d and Supplementary Video 2), most iCasp9 cells treated with 0.25 nM inducer died with high FRET Ratios (i.e. the FRET reporter was cleaved by activated Caspase3, Supplementary Fig. 1e and Supplementary Video 3). These results indicated that the FRET Ratio indeed is a good reporter for cell fate. To have the potential to scale up, we further validated that the FRET reporter can also report cell fate accurately through flow cytometry measurements (Supplementary Fig. 2a).

Using the iCasp9 cell system above, we characterized heterogeneous cell responses to the inducer. We focused on 0.25 nM inducer first as this dose provided comparable population sizes of survival and dead cells in our experimental system. We treated iCasp9 cells with 0.25 nM inducer and tracked them using a confocal microscope for 24 h as apoptotic events had largely ceased by this time (Fig. 1d). Longer treatment time would result in a decreased death percentage due to the division of survival cells (Supplementary Fig. 2b and Supplementary Video 2). We then extracted single-cell profiles of iCasp9-mCherry (Fig. 1e) and FRET Ratio (Fig. 1f) from surviving and dead cells, respectively. A significant cell-to-cell variability was observed. Some cells were dead and characterized with a significant decrease of iCasp9-mCherry and a sharp increase of FRET Ratio (pink lines, Fig. 1e,f), while some cells survived with various iCasp9-mCherry profiles but a constantly low FRET Ratio (blue lines, Fig. 1e,f).

### Heterogenous cell fates dictated by initial iCasp9 level and XIAP/C3 ratio

To investigate the source of heterogenous cell fates, we compared the profiles of both iCasp9-mCherry and FRET Ratio in survival cells (Fig. 2a) to those in dead cells (Fig. 2b). As expected, the survival cells showed a constantly low FRET Ratio, while the dead cells exhibited a sharp increase of the FRET Ratio. Unexpectedly, the dead cells showed a significant higher initial iCasp9-mCherry level than that in survival cells (Fig. 2c). We further binned the cells by the initial iCasp9 level, and found a clear dependency of the death percentage on the initial iCasp9 level (Fig. 2d), suggesting the cell fate is highly correlated with its initial iCasp9 level.

**Fig. 2:**
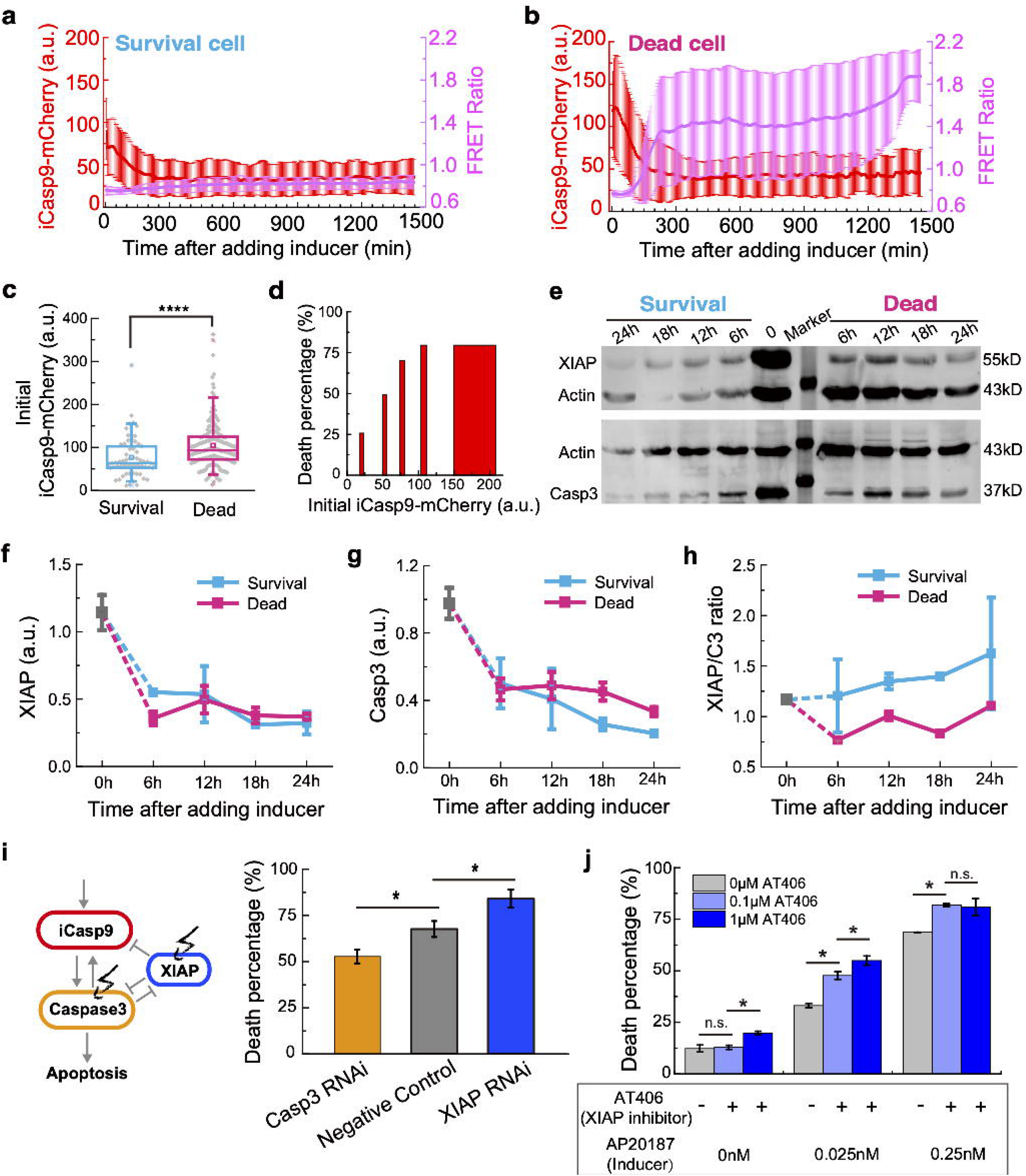
Initial iCasp9 level and XIAP/C3 ratio in survival and dead cells. **a and b,** The averaged iCasp9-mCherry fluorescence signal (shown in red) and FRET Ratio signal (shown in purple) in survival cells (**a**) and dead cells (**b**) after treatment of 0.25 nM inducer. The data for survival cells are averaged from 40 independent measurements, and the data for dead cells are averaged from 68 independent measurements. Error bars represent ± standard deviation. **c,** The initial iCasp9-mCherry level in survival and dead cell populations treated with 0.25 nM inducer. The boxes represent the interquartile range between the first and third quartiles, whereas the whiskers represent the 95% and 5% values, and the squares represent the average. **d,** Cell death percentage is plotted against the initial iCasp9-mCherry expression level. **e,** Western Blot for XIAP (upper panel) and Caspase3 (lower panel) in survival and dead cells after treating with 0.25 nM inducer for 6 h, 12 h, 18 h, and 24 h. Actin was used as a loading control. **f-h,** Quantitative analysis of XIAP level (**f**), Caspase3 level (**g**), and XIAP/C3 ratio (**h**) in survival cells (blue) and dead cells (pink) in different time points. **i,** Perturbations of XIAP or Caspase3 expression level by siRNA. Death percentage for cells treated with XIAP siRNA (blue) or Caspase3 siRNA (orange) is compared with a negative control (grey). **j,** Death percentage for cells treated with different concentrations of an XIAP inhibitor AT406 in combination with 0.025 nM inducer (light blue), 0.25 nM inducer (blue) and no inducer (grey). Error bars represent ± standard deviation. * *P* < 0.05; **** *P* < 0.0001.

However, the death percentage reached a plateau of ~80% when the initial iCasp9-mCherry level increased to 100 (Fig. 2d). In addition, the initial iCasp9 level in survival and dead cells had a big overlap, though the difference between them was significant (P<0.0001) (Fig. 2c). All these results suggested that the initial iCasp9 level is not the sole determinant of the cell fate, which led us to consider the potential effect of the other two players in AP20187-induced apoptosis pathway: XIAP and Caspase3. Thus, we measured their amount in both survival and dead cells treated with 0.25 nM inducer for different time (Fig. 2e). Compared with the initial values (0 h), both XIAP and Caspase3 dramatically decreased after adding inducer for 6 h, even in the survival cells (Fig. 2f,g), indicating a fighting process between XIAP and Caspase3^31^. As we further quantitatively compared the amount of XIAP and Caspase3 between dead and survival cells, a significant difference on XIAP/C3 ratio was observed at all time points (Fig. 2f-h). The XIAP/C3 ratios in survival cells were significantly higher than that in dead cells at all time points (Fig. 2h), suggesting that cell survives due to the overwhelming XIAP to Caspase3 level.

To further confirm the effect of XIAP/C3 ratio on cell fate, we knocked down the level of either XIAP or Caspase3 by RNAi. With a knockdown efficiency of 68% on XIAP, the death percentage was increased from 68% to 84%, while a knockdown efficiency of 59% on Caspase3 led to a decrease of death percentage from 68% to 53% (Fig. 2i). These results indicate that the variability of XIAP and Caspase3 levels in single cells also contributes to the cell fate heterogeneity. Given the numerous XIAP-targeted drugs, these results opened up a potential therapeutic strategy of combining iCasp9 inducer with XIAP inhibitors to achieve a higher killing efficiency. We tested this strategy in our system, using AT406, a small molecule drug, for XIAP inhibition. Indeed, a significant increase of killing efficiency was observed in all inducer concentrations (Fig. 2j).

Taken together, by employing the iCasp9 cell system we constructed, we found that the cell-to-cell variability in the initial iCasp9 level and the XIAP/C3 ratio are the main two contributors for the cell fate heterogeneity.

### Different types of survival cells

In addition to the heterogenous survival and dead cell fates, substantial heterogeneity was also found within both the survival and dead cell populations. As for the survival cells, we observed several distinct types in light of their iCasp9-mCherry and FRET Ratio profiles, indicating that they survive because of different reasons.

Based on whether there was a decrease in iCasp9-mCherry signal after adding the inducer, survival cells are divided into two groups. Those with no decrease of iCasp9-mCherry (< 50%) were grouped as survival cell type 1, indicating an insufficient induction of iCasp9 (red lines, Fig. 3a). All type 1 cells showed constantly low FRET Ratio signals (purple lines, Fig. 3a). The insufficient iCasp9 induction was presumably due to either low initial iCasp9 level or low inducer uptake efficiency. Thus, we further classified type 1 survival cells into two sub-populations according to their initial iCasp9-mCherry levels. Since the dead probability of cells with initial iCasp9-mCherry level above 60 started to be higher than the survival probability (Supplementary Fig. 3), we grouped cells with initial iCasp9-mCherry level below 60 as survival cell type1-a (left panel, Fig. 3a) while the remaining cells as survival cell type1-b (right panel, Fig. 3a).

**Fig. 3:**
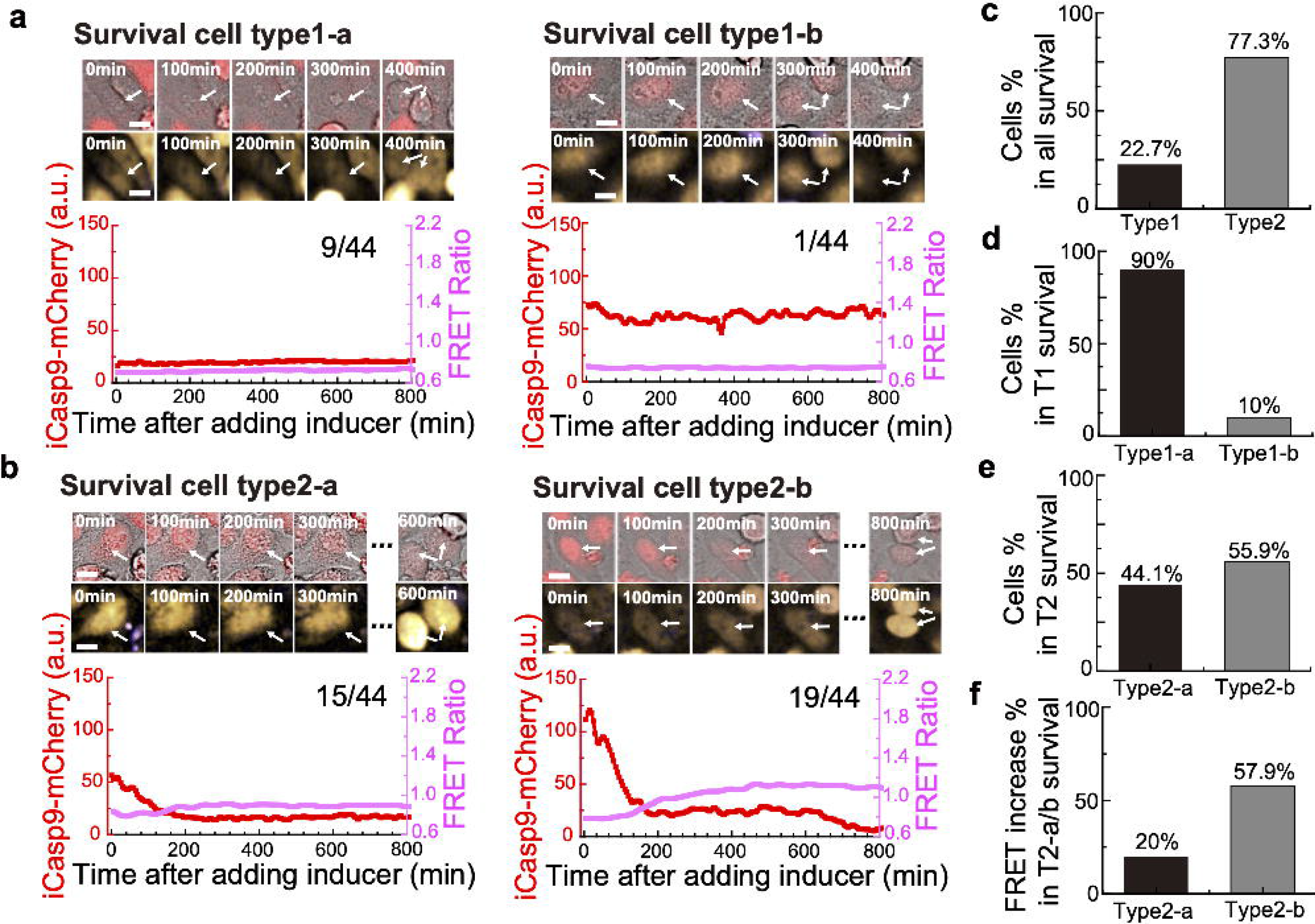
Substantial heterogeneity exists within survival cell population. **a and b,** Different types of cells surviving from the treatment of inducer. The upper panels show representative images (BF and iCasp9-mCherry merge on top, CFP and FRET merge on bottom) of different survival cell types. The lower panels show the trajectory of iCasp9 dimerization (red) and Caspase3 activation (purple) measured from the fluorescence images above. (**a,** left) A survival cell with low initial iCasp9-mCherry level and no iCasp9-mCherry drop, and a constantly low FRET Ratio. (**a,** right) A survival cell with high initial iCasp9-mCherry level and no iCasp9-mCherry drop, and a constantly low FRET Ratio. (**b,** left) A survival cell with low initial iCasp9-mCherry level and an iCasp9-mCherry drop, and a constantly low FRET Ratio. (**b,** right) A survival cell with high initial iCasp9-mcherry level and an iCasp9-mCherry drop, and a slight increase of FRET Ratio. **c,** The percentage of type 1 and type 2 survival cells in all survival cells studied. **d**, The percentage of type1-a and type1-b survival cells in type1 survival cells. **e,** The percentage of type2-a and type2-b survival cells in type 2 survival cells. **f,** The percentage of survival cells with FRET Ratio increase by at least 15% of the initial ratio in type2-a and type2-b survival cells.

For those survival cells with significant decrease of iCasp9-mCherry, we grouped them into survival cell type 2 (Fig. 3b). In contrast to the constantly low FRET Ratio in survival cell type 1, we observed a slight increase of FRET Ratio happening concomitantly with the decrease of iCasp9-mCherry in survival cell type 2, suggesting that Caspase3 had been partially activated. The final surviving cell fate was reached in these cells presumably due to a high XIAP level overwhelming the apoptotic function of iCasp9 and Caspase3. Based on whether the initial iCasp9 level was higher than 60, we also classified type2 cells into two sub-populations, survival cell type2-a (initial iCasp9<60, left panel, Fig. 3b) and survival cell type2-b (initial iCasp9>=60, right panel, Fig. 3b), respectively.

To investigate the contributions of each survival mode, we further quantified the number of survival cells from different types. Among 44 survival cells we analyzed under the condition of 0.25 nM inducer, only 22.7% (10/44) of cells had no obvious iCasp9 dimerization (Fig. 3c, Type 1), and 90% (9/10) of them showed the initial iCasp9 level lower than 60 (Fig. 3d, Type1-a), indicating the importance of initial iCasp9 level for an effective iCasp9 dimerization. The remaining cells (1/10) showed reasonably high initial iCasp9 level but no sign of dimerization, suggesting a low efficiency of drug uptake (Fig. 3d, Type1-b)^32^. However, 77.3% (34/44) of cells presented significant iCasp9 drops (Fig. 3c, Type 2), implying that most of the survival cells had an effective iCasp9 dimerization. Among this population, 44.1% (Fig. 3e, Type2-a) and 55.9% (Fig. 3e, Type2-b) of the cells had initial iCasp9 level lower and higher than 60, respectively. Though all Type 2 cells survived from the fighting between XIAP and Caspases, fewer cells in Type2-a had an increase of FRET Ratio (> 15%) than that in Type2-b (Fig. 3f), indicating higher initial iCasp9 level would generally leads to higher Caspase3 activation.

### Substantial heterogeneity exists within dead cell population

Then we looked into the dead cell population. Though a significant drop of iCasp9-mCherry fluorescence and a sharp increase of FRET Ratio were observed in all dead cells (Fig. 4a), the dynamics of the iCasp9 dimerization (Fig. 4b) and Caspase3 activation (Fig. 4c) showed a large cell-to-cell variability.

**Fig. 4:**
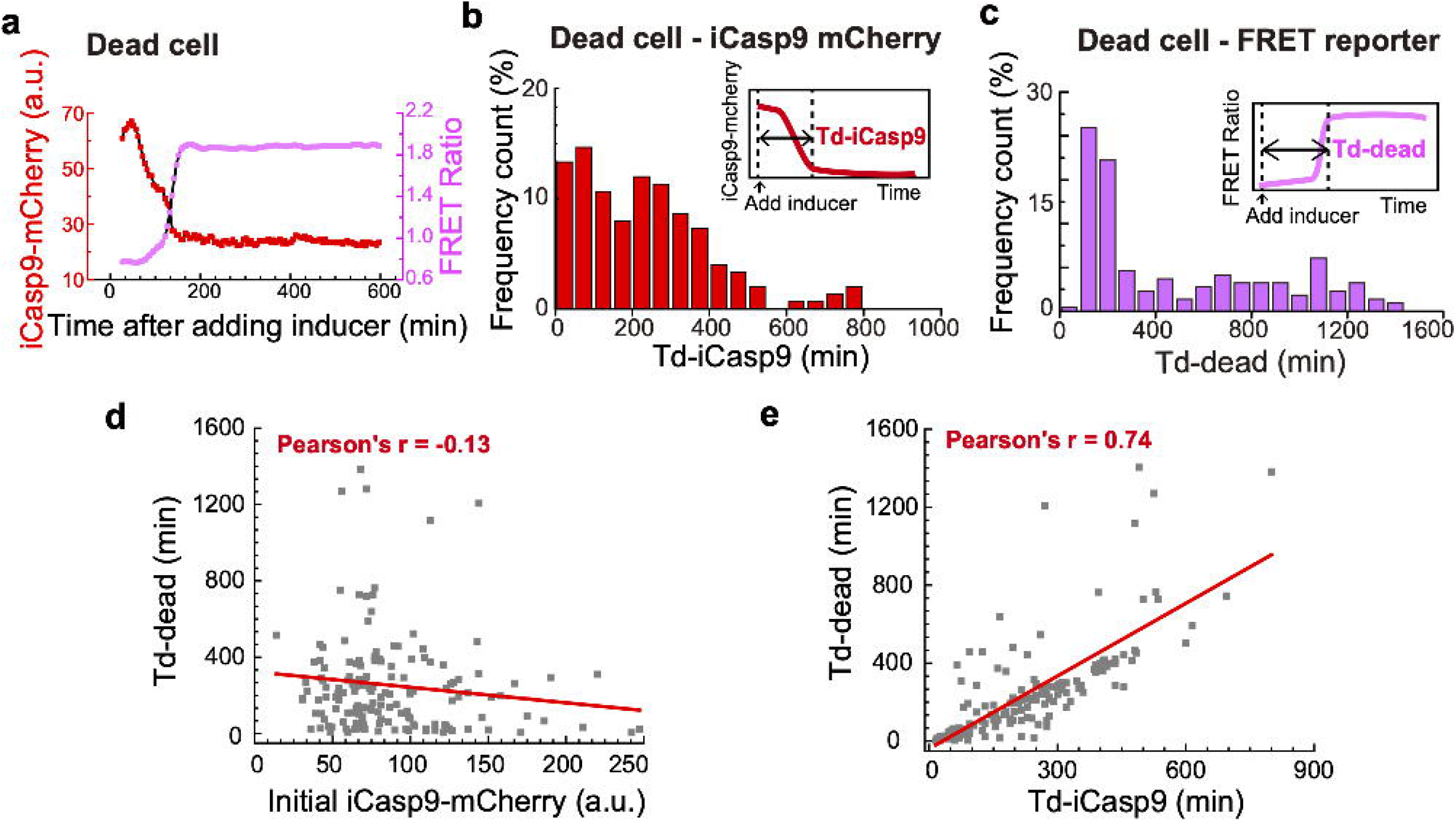
Cell-to-cell variability of death timing in dead cell population. **a,** Fluorescence signals of iCasp9-mCherry and FRET ratio in a typical dead cell. **b,** Frequency count of dead cells with different iCasp9 dimerization time under the condition of 0.25 nM inducer. Inner image represents a schematic view for iCasp9 dimerization time ‘Td-iCasp9’, which is defined as the time duration from adding inducer to the iCasp9-mCherry fluorescence signal reaching 10% higher than the minimum. **c,** Frequency count of dead cells with different death time under the condition of 0.25 nM inducer. Inner image represents a schematic view for cell death time ‘Td-dead’, which is defined as the time duration from adding inducer to the FRET ratio reaching 90% of the maximum. **d**, Td-dead is plotted against the initial iCasp9-mCherry level in dead cells. Pearson’s correlation is −0.13. **e,** Td-dead is plotted against Td-iCasp9 in dead cells. Pearson’s correlation is 0.74.

To quantify the heterogeneity, we defined “Td-iCasp9” as the time for iCasp9-mCherry intensity decreasing from the initial level to the level at 10% higher than the minimum (Fig. 4b), and “Td-dead” as the time from adding inducer to the FRET Ratio reaching 90% of its maximum (Fig. 4c), respectively.

As iCasp9 dimerization was the effective input of the circuit, we asked what features of iCasp9 determines the variability of death timing. We first looked at the initial iCasp9 level, which is correlated with cell survival/death fates. However, only a weak negative correlation (Pearson’s r=−0.13) was found between Td-dead and initial iCasp9 level (Fig. 4d). Then we analyzed the correlation between Td-dead and Td-iCasp9 (Fig. 4e). A strong positive correlation (Pearson’s r=0.74) was seen, indicating cells with faster iCasp9 dimerization are more likely to die faster than cells with slower iCasp9 dimerization. Different dynamics of iCasp9 dimerization was presumably due to the cell-to-cell variability on drug transportation efficiency^32^.

### Dose-effect of inducer on caspase dynamics and cell fates

Since drug dose is the easiest thing to be adjusted in clinic, we further varied the inducer concentration and investigated the dose-effect on cellular responses. We first systematically investigated the killing efficiency of different inducer concentrations, ranging from 0.0001 nM to 100 nM (Fig. 5a). While the wild-type cells which do not incorporate iCasp9 gene did not respond to the inducer (black dots, Fig. 5a), iCasp9 cells showed a significant enhancement of cell death with increasing inducer concentration (red dots, Fig. 5a). To understand the dose-effect of inducer at single cell level, we further looked into the dynamics of iCasp9 dimerization and Caspase3 activation. Here, we took 0.025 nM, 0.25 nM and 2.5 nM as representatives of the low, medium and high concentrations, given the cell death percentages in these three concentrations were ~30%, ~60% and ~90%, respectively (blue, green and orange dash lines, Fig. 5a).

**Fig. 5:**
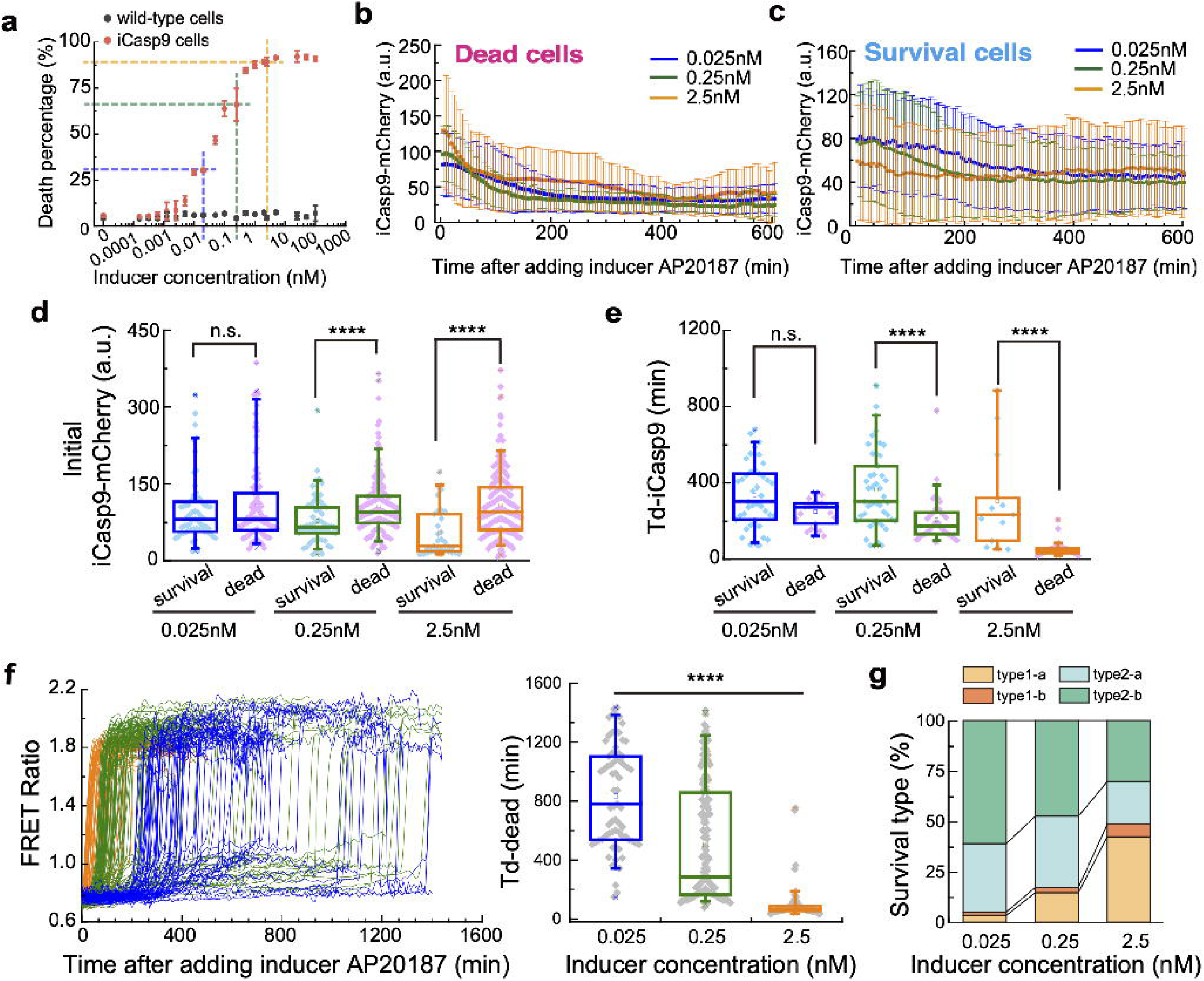
Inducer dosage affects caspase dynamics, cell-to-cell variability and cell death. **a,** Cell death percentage is plotted against different concentrations of inducer. Data are shown as mean ± standard deviation. **b and c,** Averaged trajectories of iCasp9-mCherry for dead cells (**b**) and survival cells (**c**) treated with 2.5 nM (orange), 0.25 nM (green) and 0.025 nM (dark blue) inducer. Error bars represent ± standard deviation. **d and e,** Initial iCasp9-mCherry level (**d**) and Td-iCasp9 (**e**) in survival (blue dots) and dead (pink dots) cells after the treatment of 0.025 nM, 0.25 nM and 2.5 nM inducer. **f,** The trajectories of FRET Ratio for individual dead cells in the condition of 0.025 nM, 0.25 nM and 2.5 nM inducer (**f**, left panel). Td-dead for dead cells treated with 0.025 nM, 0.25 nM and 2.5 nM inducer (**f**, right panel). **g,** Percentage of four survival cases in cell population treated with 0.025 nM, 0.25 nM and 2.5 nM inducer. The boxes of the box plot represent the interquartile range between the first and third quartiles, whereas the whiskers represent the 95% and 5% values, and the squares represent the average. * *P* < 0.05; ** *P* < 0.01, *** *P* < 0.001, **** *P* < 0.0001.

We firstly extracted the average trajectories of iCasp9-mCherry from all dead (Fig. 5b) and survival cells (Fig. 5c). Obvious differences were found both between dead and survival cells and among different concentrations. The survival cells showed a significantly lower initial iCasp9 level than that in dead cells under concentrations of 0.25 nM and 2.5 nM, further confirming our conclusion that cells with higher initial iCasp9 level are more easily to die (Fig. 5d). No significant difference on initial iCasp9 level was found between dead and survival cells in the concentration of 0.025 nM, presumably due to the extremely low killing efficiency at that concentration, which was just slightly higher than the background death percentage (Fig. 5a). In this case, the contribution to cell fate from other factors, such as the XIAP and Caspase3 level, may shadow that of the initial iCasp9 level. It is worth to note that a decreasing initial iCasp9 level was found in survival cells as the inducer concentration increases, indicating that some survival cells with high initial iCasp9 level can be killed by increasing the inducer concentration (Blue dots, Fig. 5d).

As for the dynamics of iCasp9 dimerization, we measured the Td-iCasp9 for survival and dead cells under different inducer concentrations. For survival cells, since there was no obvious iCasp9 dimerization in type1 cells, we only extracted the averaged trajectories of iCasp9-mCherry from type 2 cells for Td-iCasp9 analysis (Supplementary Fig. 4). It turned out that, even type 2 cells had iCasp9 dimerization, their Td-iCasp9 was significantly longer than that in dead cells, suggesting that survival cells may generally have a weak capability in drug uptake. Moreover, both survival and dead cells possessed a shortened Td-iCasp9 when being treated with higher concentrations, suggesting higher inducer concentration would speed up the iCasp9 dimerization.

To further support our findings that faster iCasp9 dimerization can accelerate cell death, we extracted the trajectories of FRET Ratio in each dead cell under different inducer concentrations (left panel, Fig. 5f). Quantitative analysis on Td-dead (right panel, Fig. 5f) showed that cells died at an averaged time of ~800 min after being treated with 0.025 nM inducer, with the 5^th^ and 95^th^ percentile values of 300 min and 1400 min. However, in the condition of 2.5 nM inducer, the averaged Td-dead was dramatically shortened to ~80 min and the variability was also greatly reduced. These results led us to propose that slow dimerization of iCasp9 resulted in a gradual accumulation of active iCasp9, which might be easily neutralized by XIAP and thus hampers cell death. We further quantified Td-iCasp9 and cell death percentage in different inducer concentrations. The results showed cell death percentage was negatively related to Td-iCasp9, confirming that rapid iCasp9 dimerization is more potent to causing cell death (Supplementary Fig. 5).

Besides the impact on death timing in dead cells, drug dose also changed the composition of survival cell types. We found the survival cells with low initial iCasp9 level became dominant in higher concentrations (type1a+type2a, Fig. 5g).

Taken together, higher concentration of AP20187 would speed up the iCasp9 dimerization, and lead to faster death with lower cell-to-cell variability.

### Development of drug resistance in multiples rounds of inducer treatmenet

Substantial heterogeneity of responsiveness has been shown in clonal iCasp9 cell population, and that higher concentration could lead to faster death with lower cell-to-cell variability. However, since there were always some cells left unkilled in the drug treatment, we wonder if a complete or near complete killing could be achieved by applying multiple rounds of treatment. We exposed iCasp9 cells to multi-rounds of 0.25 nM inducer treatment (Fig. 6a). After the first round of treatment, survival cells remained attached to the dish whereas dead cells detached, allowing us to recover survival cells by trypsinization. We plated survival cells into a fresh medium without inducer and cultured them for 48 h. Then these survival cells were challenged with a second dose of 0.25 nM inducer. We cultured cells survived from the second dose for another 48 h and treated them with a third dose of 0.25 nM inducer. For each round of inducer treatment, cells were treated with inducer for 24 h and the death percentage was monitored (Fig. 6b). Instead of approaching higher killing efficiency, an obvious accumulation of drug resistance was observed with repeated rounds of treatment. While 65% of cells died after the first round of treatment, the death percentage dropped to 48% for the second round and only 12.3% of cells were killed in the 3^rd^ round treatment (Fig. 6b).

**Fig. 6:**
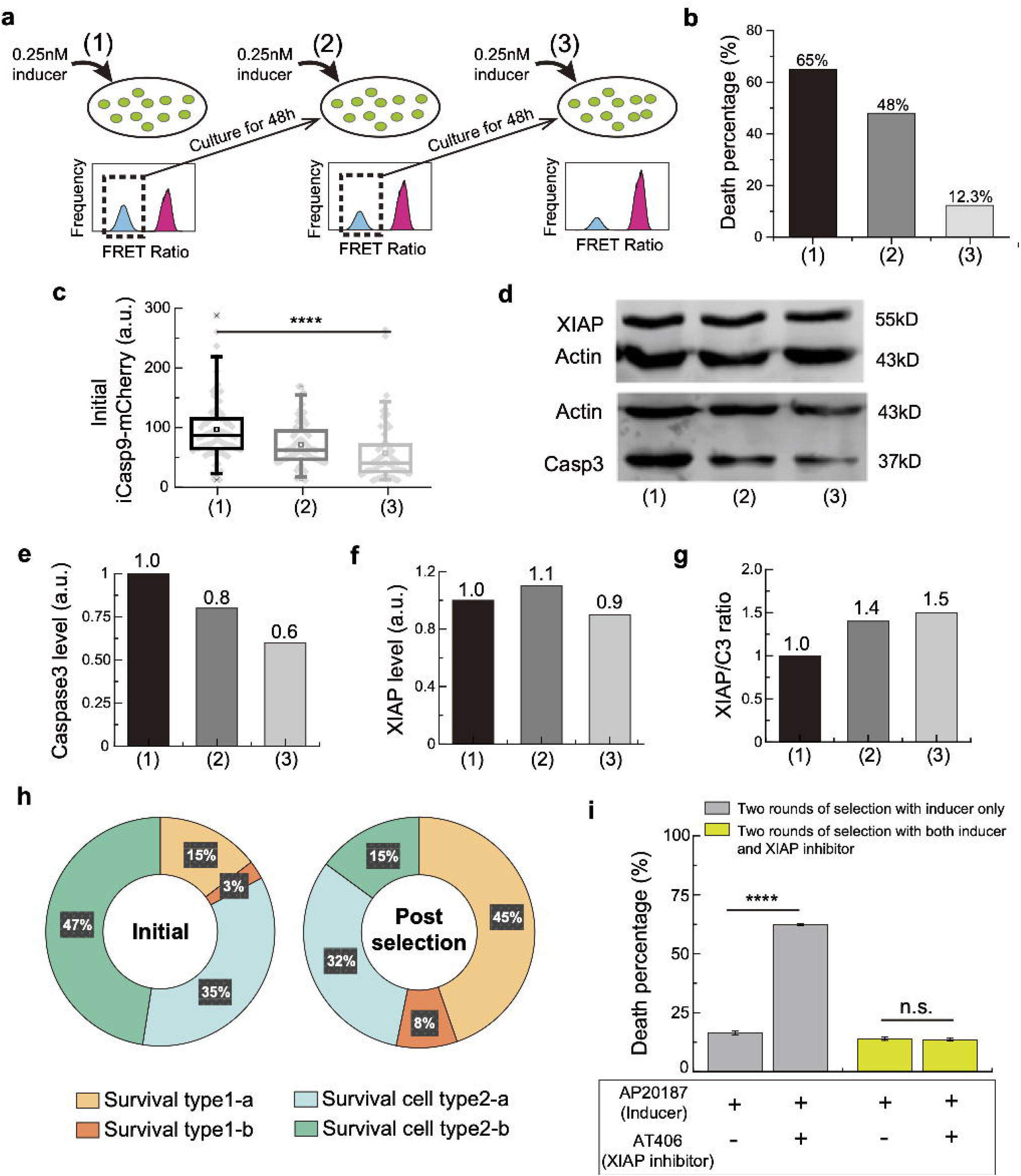
Multiple rounds of inducer treatment place selective pressure on protein level and lead to drug resistance. **a,** Schematic view of the multi-drugging experiment. Survival cell population with low FRET Ratio (shown in blue) is collected and re-cultured for two days before the next round of drugging. **b,** Death percentage for cells after each round of treatment with 0.25 nM inducer. **c,** Initial iCasp9-mCherry expression in cells before each round of inducer treatment. The boxes represent the interquartile range between the first and third quartiles, whereas the whiskers represent the 95% and 5% values, and the squares represent the average. **d,** Western Blot for XIAP (upper panel) and Caspase3 (lower panel) in cells before the first (column 1), the second (column 2), and the third (column 3) round of inducer treatment. Actin was used as a loading control. **e-g,** Quantitative analysis of Caspase3 expression (**e**), XIAP expression (**f**), and XIAP/C3 ratio (**g**) in cells before each round of inducer treatment. **h,** Percentage of four survival cell types after the first treatment (left pie chart) and after the third treatment (right pie chart) of the inducer. **i,** Death percentage of post-selection cells at the third round of treatment with either the inducer only or the combination of the inducer and the XIAP inhibitor. The cells were selected by two rounds of treatment with 0.25 nM inducer only (grey bars) or with a combination of 0.25 nM inducer and 0.1 μM XIAP inhibitor AT406 (yellow-green bars). Data are shown as mean ± standard deviation.

To further investigate the origin of the accumulating drug resistance, we quantified the iCasp9, XIAP and Caspase3 protein levels in cells right before each round of inducer treatment. A significant drop of the initial iCasp9-mCherry level was observed as the round increases, indicating a strong selective pressure on the initial iCasp9-mCherry level (Fig. 6c). Meanwhile, we also found a decrease of Caspase3 level, while XIAP maintained at a similar level (Fig. 6d-f). As a result, the XIAP/C3 ratio was higher in cells survived from multiple rounds of treatment (Fig. 6g).

As discussed before, cells survive from the inducer treatment in different ways (Fig. 3). We compared the composition of different survival cell types after the first and the third round of treatment (Fig. 6h). While there was a total of 50% (type1a+type2a) survival cells (initial iCasp9 < 60) after the first treatment (left pie chart, Fig. 6h), the percentage increased to 77% after three rounds of treatment (right pie chart, Fig. 6h). These results further support the selective pressure on the initial iCasp9 level. Interestingly, the percentage of survival cells due to high XIAP/C3 ratio (type2-b) dropped dramatically from 47% to 15%, suggesting minor contributions of XIAP/C3 ratio to the increasing drug resistance, consistent with the fact that no obvious selective pressure was observed on XIAP level (Fig. 6f). Taken together, our experiments showed that multiple rounds of treatment cannot increase the killing efficiency. Instead, it would place selective pressure on protein levels, especially on the level of initial iCasp9, leading to drug resistance.

Since no obvious change on XIAP level was found after multiple rounds of treatment, we wonder if higher killing efficiency could be achieved by combining inducer with XIAP inhibitor. Indeed, after two rounds of AP20187 treatment, by combining with 0.1 µM XIAP inhibitor AT406 at the third round, a significant increase of killing efficiency was obtained, from 16.5% to 62.4% (grey bars, Fig. 6i). Surprisingly, if the drug combination was used from the beginning, i.e., if the previous two rounds of treatment were by the combination of inducer AP20187 and XIAP inhibitor AT406, the killing efficiency at the third round was much lower with or without the drug combination (yellow-green bars, Fig. 6i). This suggests that the continuous application of combinational drugging placed selective pressure on both iCasp9 and XIAP, resulting in a dramatic attenuation of the efficacy of the XIAP inhibitor. Our findings hints a “smart strategy” of drug combination usage.

## DISCUSSION

In this paper, we constructed an iCaps9 system in which we can track the dynamics of iCasp9 dimerization and Caspase3 activation, and the cell fate simultaneously in single cells using live-cell imaging. Large heterogeneity in the clonal population of iCasp9 cells was observed. The heterogeneity not only manifested in the final cell fate of life and death, but also in the different surviving ways within survival cells and the varied death timing within the dead cells.

The major source of heterogeneous cell responses to the inducer is the cell-to-cell variability in the initial iCasp9 level and XIAP/C3 ratio. We found that there was significantly higher initial iCasp9 level and lower XIAP/Caspase3 ratio in survival cells than that in dead cells, which suggests the survival or death cell fate is mainly determined by the fighting process between the pro-apoptotic proteins and the anti-apoptotic proteins. This conclusion was further confirmed by RNAi experiments and our observations on the different survival cell types. By combining the analysis of iCasp9-mCherry and FRET Ratio profiles in single cells, we found that most of the cells (77.2%, type2 survival cells) survived through XIAP overwhelming the apoptotic effects of iCasp9 and Caspase3. Besides the variability in iCasp9, XIAP and Caspase3 levels, we also observed some influence from the inducer uptake efficiency on the heterogeneous cell response. There was a small population of cells (22.8%) that survived due to insufficient induction of iCasp9 dimerization, among which the low initial iCasp9 level was the main cause (90%) while about 10% was likely due to the low inducer uptake efficiency.

Our analysis indicated that the killing efficiency could be adjusted by changing the protein expression level. Indeed, the killing efficiency was increased with a knockdown on XIAP, while a knockdown on Caspase3 led to a decrease of the death percentage. These results suggest a big difference on the killing efficiency for various cell types or tissues/organs, as their protein expressions including Caspase3, Caspase9 and XIAP varies a lot (seen in database ‘The Human Protein Atlas’). Thus, personalized treatment with respect to specific cell types or tissues may be needed to reach the desired killing efficiency.

By looking into the dead cell population, we found a substantial heterogeneity in death timing, which was poorly correlated with initial iCasp9 level but highly correlated with the speed of iCasp9 dimerization. The heterogeneity here is most likely originated from the cell-to-cell variability in drug uptake efficiency. As the inducer AP20187 is a small molecule that can be readily diffused into cells, its cellular concentration presumably depends on the drug exclusion process^32^ which could be variable from cell to cell. To enhance the uptake efficiency, we increased the inducer concentration as this is the most feasible strategy in clinical treatment. Higher dose of inducer indeed triggered faster dimerization of iCasp9, leading to more rapid activation of Caspase3 and faster cell death with less variability. In contrast, when a low dose of AP20187 is applied, it takes longer for iCasp9 to dimerize. The slow accumulation of active iCasp9 would be more easily neutralized by XIAP. In this scenario, lower dose of AP20187 tends to cause bigger cell-to-cell variability as the slow time scale for iCasp9 activation would amplify the opposing effects of XIAP and Caspase3 expression.

Multiple rounds of inducer application did not kill more cells. Instead, the surviving cells developed heritable drug resistance. We found that the resistance came from the low iCasp9 expression. This heritable protein variation probably originated from the epigenetic differences^33^ in individual cells. Further investigation of the epigenetic source may improve iCasp9-mediated killing efficiency. Interestingly, little selective pressure on XIAP expression level was observed in the multi-round inducer treatment. Thus, we can significantly improve killing efficiency on the inducer resistant cell population by combining XIAP-targeted inhibitor with the inducer. It is worth to note that the efficacy of drug combination was dramatically diminished if used continuously from the very beginning.

The clinic application of iCasp9 suicide gene is still in its infancy. More investigations on the system are required to unleash its full potential. Our work unveils the sources of cell-to-cell variability and drug resistance in clonal population of iCasp9 cells, paving the way for improving therapeutic strategies to achieve optimal performance.

## MATERIALS AND METHODS

### Cell lines and cell culture

The human embryonic kidney 293T cells and Hela cervical cancer cell lines were gifts from Prof. Jianguo Chen. Cells were cultured in Dulbecco’s Modified Eagle’s Medium (DMEM) (Gibco, Cat# 11965092) containing 10% fetal bovine serum (Gibco, Cat# 16000044) and 100 U/mL penicillin and streptomycin (Invitrogen, Cat# 15140163) in a humidified environment with 5% CO_2_ at 37 °C.

### DNA constructs

Lentivirus vectors pHR-mCherry, pCMVdR8.91, and pMD2.G were kindly provided by Prof. Ping Wei. Full-length Caspase9 was a gift from Prof. Xiaodong Wang. The FRET reporter (pECFP-DEVDR-Venus, Cat # 24537) was purchased from Addgene and the iDimerize™ Inducible Homodimer System (Cat # 635068) was purchased from Clontech Laboratories, Inc. iCasp9 construct was built by inserting full-length Caspase9 to pHom-1 vector with a linker sequence ‘RPPPR’, which was verified by sequencing with primers Seq_Casp9_F and Seq_Casp9_R. Then the integrated DmrB-linker-caspase-9 sequence in the pHom-1 vector was cloned into the pHR-mCherry vector. We used Primer SpeI-Casp9_F and BamHI-Casp9_R to generate full-length Caspase9 fragment with SpeI and BamHI restriction enzyme cleavage site, so that we can insert it into the pHom-1 vector which has the same cleavage site. Primers for constructing and verifying DmrB-linker-caspase-9 are listed below.

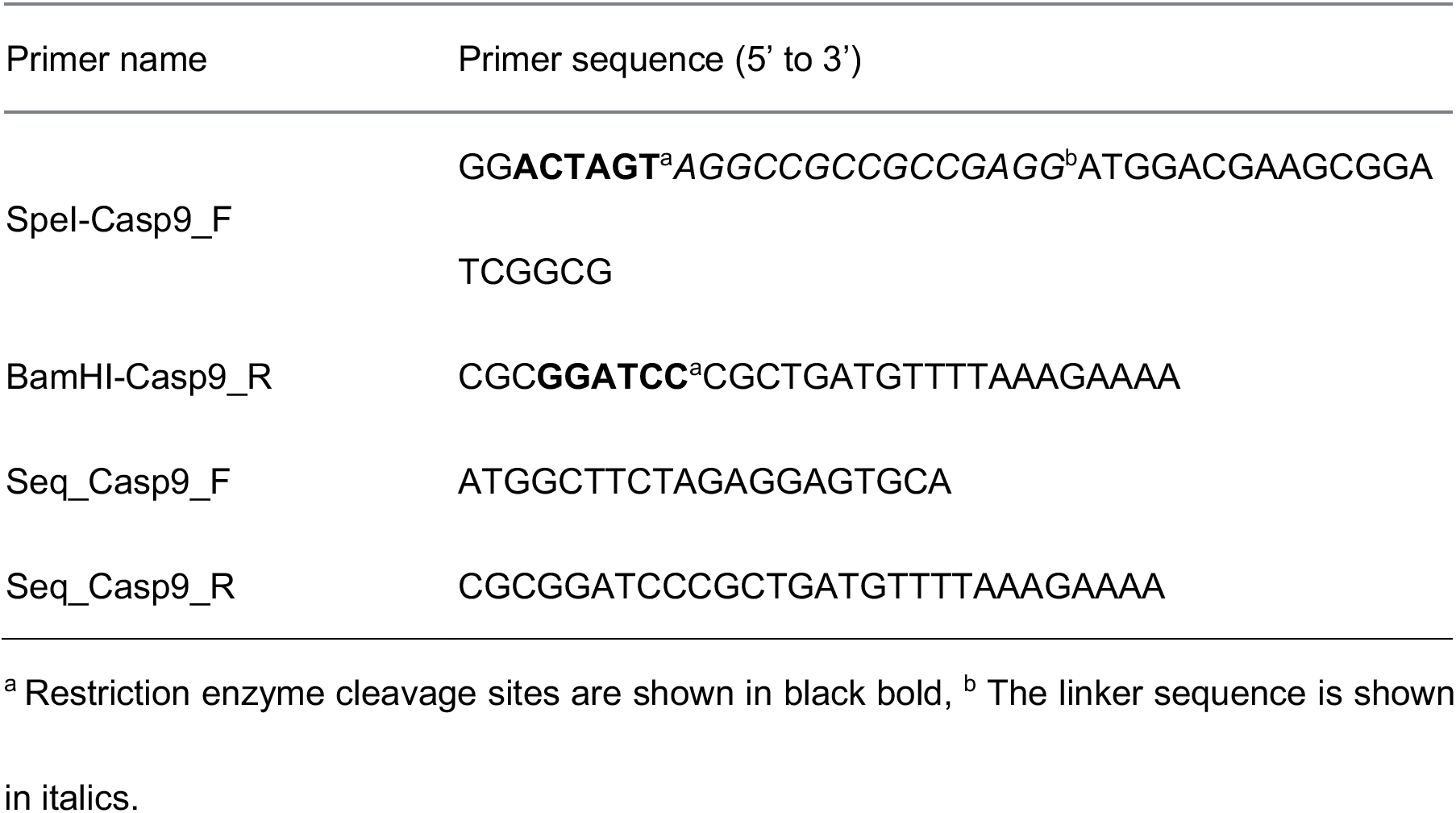

### Stable cell line construction

We transfected the FRET reporter into cells by using X-tremeGENE HP DNA Transfection Reagent (Roche, Cat# 6366236001). We mixed 1 μg of plasmid DNA (pECFP-DEVDR-Venus) and 3 μl X-tremeGENE HP DNA Transfection Reagent with 100 μl opti-MEM (Gibco, cat# 31985088), and added into cells. After transfection, we cultured the cells with DMEM medium containing antibiotic G418 (Sigma, 10131027), starting with the concentration of 1000 μg/ml and then decreasing to maintain the concentration at 200 μg/ml for ~30 days. Finally, we collected survival cells and used FACS to select and allocate single cells with positive Venus fluorescence signals into 96-well plates, and cultured every single cell into a big population as the clonal population.

Based on the stable Hela cell line with FRET reporter, the iCasp9 gene was further introduced by lentivirus infection, which was pre-generated from 293T cells. We first co-transfected three lentivirus vectors pHR-Caspase9-mCherry, pCMVdR8.91, and pMD2.G into 293T cells. After incubation in 5% CO_2_, 37°C incubator overnight, we harvested the supernatant which contains the lentivirus particles, and added them to Hela cells. Two days later, we replaced the medium with a fresh one and kept culturing cells for another week in media containing 0.5 mg/ml puromycin. Clonal population of iCasp9 cells were established as described above.

### siRNA transfection

The sequence of XIAP siRNA and Caspase3 siRNA was designed for targeting XIAP and Caspase3, respectively. NC siRNA was designed as the negative control. 100 nM siRNA was transfected into iCasp9 cells by using ROCHE X-tremeGENE™ siRNA Transfection Reagent (MilliporeSigma, Cat# 4476093001). Cells were harvested 48 hours after transfection and analyzed by western blotting.

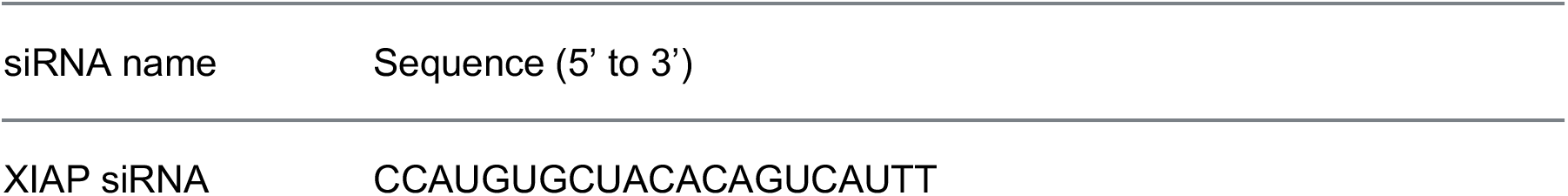
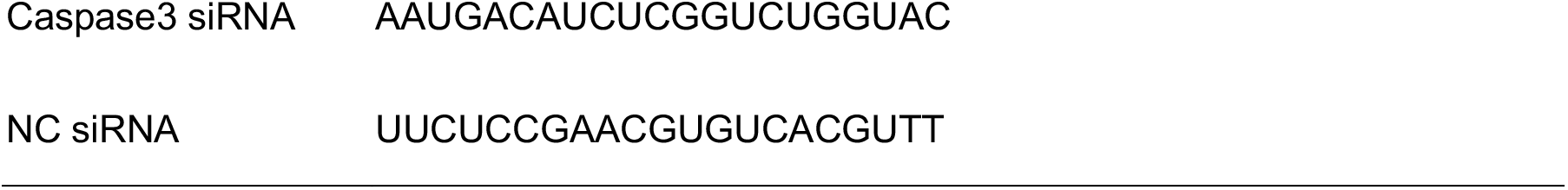

### Western blotting

For analysis of RNAi knockdown efficiency of XIAP and Caspase3, cells were harvested 48 h after transfection. For analysis of XIAP and Caspase3 at different time points, we collected survival and dead cells separately at time 6 h, 12 h, 18 h, and 24 h after the treatment of AP20187. Cells were lysed in lysis buffer (100 mM NaCl, 50 mM Tris pH 7.5, 0.5% deoxycholate, 1% Triton X-100, 0.1% SDS). Proteins were run on 12% Bis-Tris gels (cat# SL1120-500ML), then transferred to a PVDF membrane (cat# ISEQ00010) and incubated in a blocking solution (10% Skimmed Milk Powder, cat# SUP003a) 2 h at room temperature. Membranes were then incubated with the primary antibody in Can Get Signal Solution1 (cat# NKB-101) overnight at 4°C. Blots were washed three times for 5 min in washing solution (50 mM Tris-Cl pH 7.5, 150 mM NaCl, 0.05% Tween-20) and then incubated in blocking solution plus fluorescent secondary antibody in Can Get Signal Solution2 for 1 h. Membranes were washed three times in a washing solution, and we detected protein levels with fluorescent signals (LI-CORODYSSEY CLx Infrared Imaging System). Molecular weights were identified using a protein standard (BioLabs, Cat# P7712S). Beta-actin was used as the loading control. Antibodies used for western blotting are listed below:

- Primary antibody for XIAP (Cell Signaling Technology, Cat# 2042S)
- Primary antibody for Caspase3 (cat# 9662S)
- Primary antibody for Caspase9 (cat# 9502S)
- Primary antibody for Actin (cat# 3700S)
- Secondary antibody (anti-mouse cat# 926-32211 & anti-rabbit cat# 926-68070)

### Flow cytometry analysis

Flow cytometry FACS Aria (Becton-Dickenson, USA) was used to perform sorting for iCasp9 cell lines. We used a 100 μm nozzle and set the forward scatter (FSC) and side scatter (SSC) to 300 V and 240 V, respectively, to identify the population of single live cells. We set the short-RFP fluorescence parameter to 500 V, to identify the population of iCasp9-mCherry cells. We set the BV421 and BV510 fluorescence parameters to 380 V and 320 V, respectively, to identify the fluorescence signals of CFP, FRET, and Venus.

We used the high-throughput mode of flow cytometry to determine cell death percentage. The ratio of CFP to FRET was used to identify live from dead ones. Cells were plated in 96-well flat-bottom plates (Corning) with a density of 40,000 per well. Cells were then treated with AP20187 (Clontech Laboratories, Cat # 635059) in different concentrations after culturing overnight. After incubation for 24 h, cells were first centrifuged at 2000 rpm for 5 min to avoid any loss of dead cells. Then we trypsinized and collected all the cells with 200 μl medium per well of a 96-well plate as samples. FlowJo software was used to analyze death percentage based on the FRET Ratio and light scattering.

### Time-lapse microscopy

All images were captured by a Nikon Ti inverted microscope equipped with 40×/0.95 Plan Apo (Numerical aperture 1.4) objective lens and the Perfect Focus System (Nikon Co., Tokyo, Japan) for continuous maintenance of the focus. Cells were plated in 4-chamber glass-bottom dishes (In Vitro Scientific) with a density of 10,000 per well and maintained in a 37°C, 5% CO2 incubation chamber for 24 h imaging. We used filter sets that are optimized for the detection of mCherry, CFP, FRET, and Venus fluorescence. CFP fluorescence was excited at 440 nm with a 100 mW solid-state laser and collected with a 485/60 emission filter (Chroma Technology Corp). We set the exposure time at 500 ms. FRET signal was excited by 440 nm laser with an exposure time of 500 ms and collected with a 550/49 emission filter. Venus fluorescence was excited at the wavelength of 514 nm with a 100 mW solid-state laser and collected with a 550/49 emission filter. The mCherry fluorescence was excited at 561 nm with a 100 mW solid-state laser and collected with a 615/70 emission filter (Chroma Technology Corp). All images were captured at an interval of 5 min for 24 h.

### Image processing

We used ImageJ (National Institute of Health), Cellprofiler (Broad Institute), and Matlab (Mathworks) for image processing. ImageJ was used to subtract the background of images and transform the image format to adapt the software Cellprofiler. Cell segmentation and tracing were accomplished by the Cellprofiler pipeline. Firstly, cell boundary was identified based on the Venus signal to track cells at every time point. Then, we quantified the mean intensity of CFP, FRET, and Venus signals in each cell for every time point. Finally, all these measurements were exported as EXEL files for the following analysis. Further analysis was performed by the Matlab. We extracted trajectories of FRET Ratio and iCasp9-mCherry, and characterized them with the parameter of Td-Dead and Td-iCasp9.

## ACKNOWLEDGEMENTS

We thank Profs. Xiaodong Wang, Jianguo Chen and Ping Wei for kindly providing us the cells and DNA plasmids, Lucas Carey for helpful discussions, Tanqiu Liu for image analysis, and Jing Xia for data acquisition (microtubule-mCherry).

## CONFLICT OF INTERESTS

The authors declare no competing interests.

## AUTHOR CONTRIBUTIONS

C.T. and Y.Y. designed the project. Y.Y. and Y.L. designed and performed the experiments. Y.Y., H.R., and S.Q. analyzed the data. C.T. and X.Y. supervised the whole project; Y.Y., X.Y., and C.T. wrote the paper.

## FUNDING

This work was supported by The National Key Research and Development Program of China: 2018YFA0900700 and the National Natural Science Foundation of China (12090053 and 32088101).

## Data Availability Statement

All data generated or analyzed during this study are included in the main text and the supplementary information files.

## Notes

### Competing Interest Statement

The authors have declared no competing interest.

