## supplementary information for "Cell-to-cell variability in inducible Caspase9-mediated cell death"

**This PDF file includes:**

Supplementary Text
Supplementary Figs. 1 to 6

**Other Supplementary Materials for this manuscript include the following:**

Supplementary Videos 1 to 3

### SUPPLEMENTARY INFORMATION

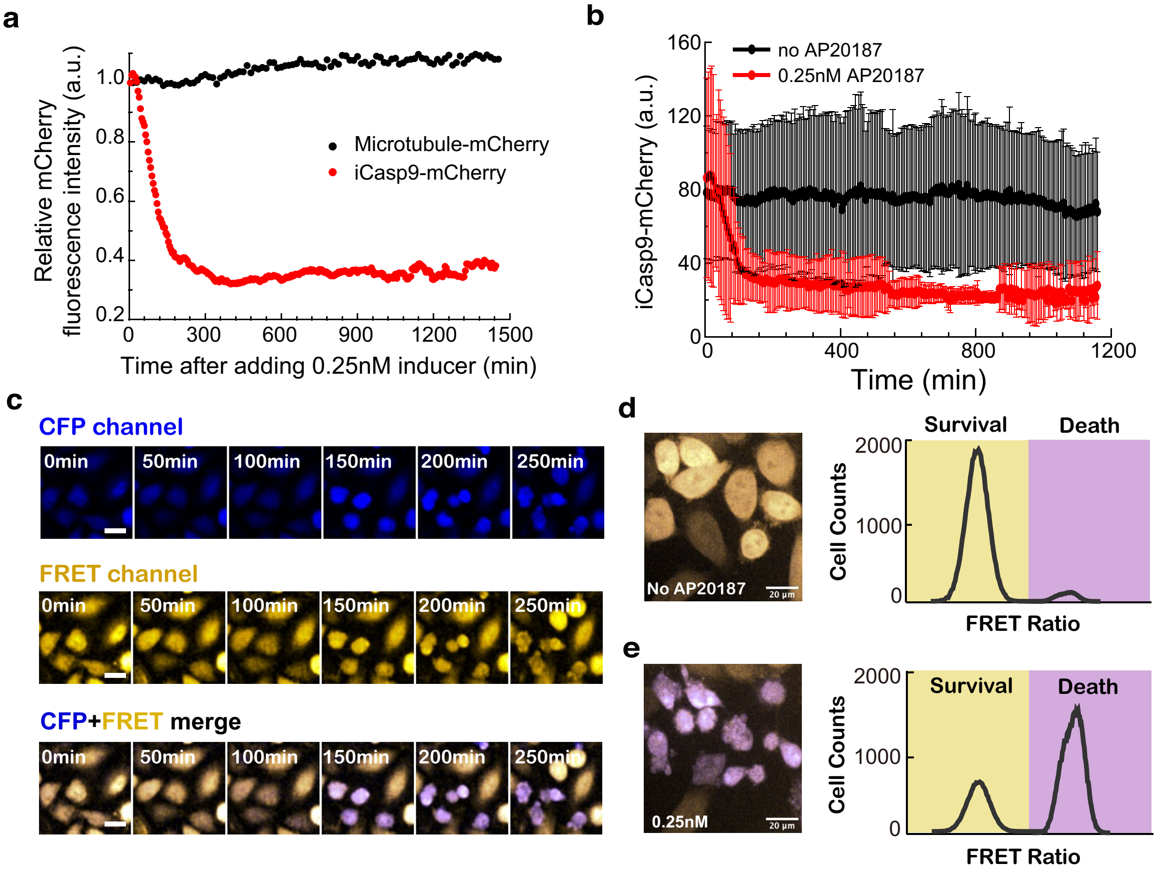


**Supplementary Fig. 1 Validation of iCasp9 cell system. a,** Fluorescence signals of iCasp9-mCherry and microtubule-mCherry in cells after adding 0.25 nM inducer AP20187. **b,** Fluorescence signals of iCasp9-mCherry in iCasp9 cells treated with and without 0.25 nM inducer AP20187. Error bars represent ± standard deviation. **c,** Time sequence images of iCasp9 cell after adding 0.25 nM inducer AP20187. Top, CFP channel; Middle, FRET channel; Bottom, CFP and FRET merge channel. FRET Ratio is defined as the ratio of CFP intensity versus FRET intensity. **d and e,** Microscopy images (left panel) and flow cytometry analysis of FRET Ratio (right panel) for iCasp9 cells without inducer AP20187 treatment (**d**), and treated with 0.25nM AP20187 (**e**). Related to Fig 1

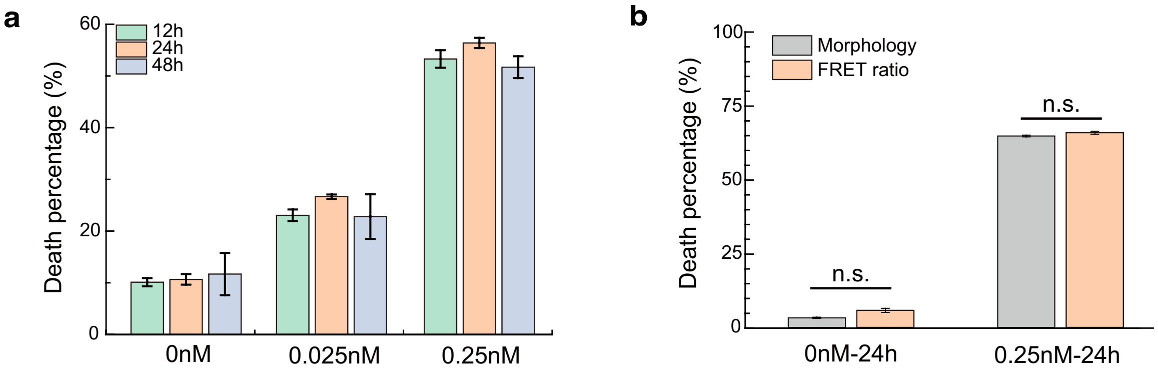


**Supplementary Fig. 2 Characterization of cell death percentage. a,** Death percentage analyzed based on cell morphology and the FRET Ratio. **b,** Death percentage of iCasp9 cells after the addition of 0 nM, 0.025 nM and 0.25 nM inducer AP20187 for 12 h, 24 h and 48 h, respectively. Data are shown as mean ± standard deviation. Related to Figure 1

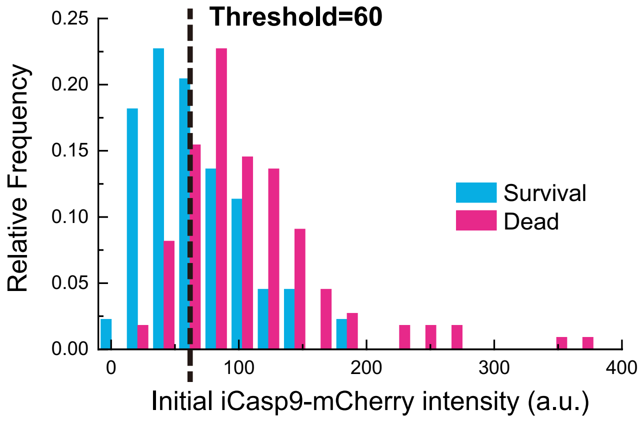


**Supplementary Fig. 3 Threshold setting for initial iCasp9 level.** Threshold of initial iCasp9 level set to distinguish survival cells with high initial iCasp9 level from those with low initial iCasp9 level. Related to Figure 3 and Figure 6

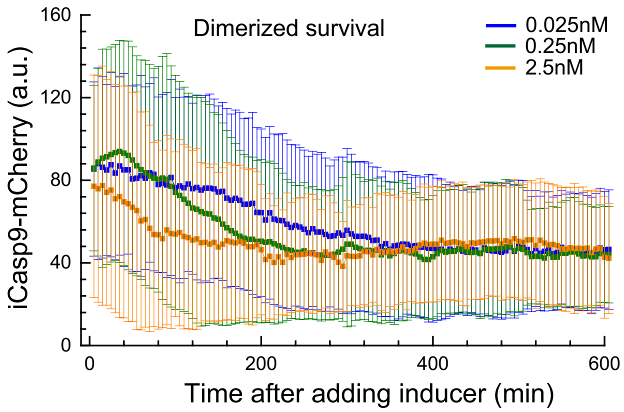


**Supplementary Fig. 4 Averaged dynamics of iCasp9-mCherry in survival cells with iCasp9 dimerization.** Averaged dynamics of iCasp9-mCherry in survival cells with iCasp9 dimerization in condition of 0.025 nM, 0.25 nM and 2.5 nM inducer. Error bars represent ± standard deviation. Related to Figure 5

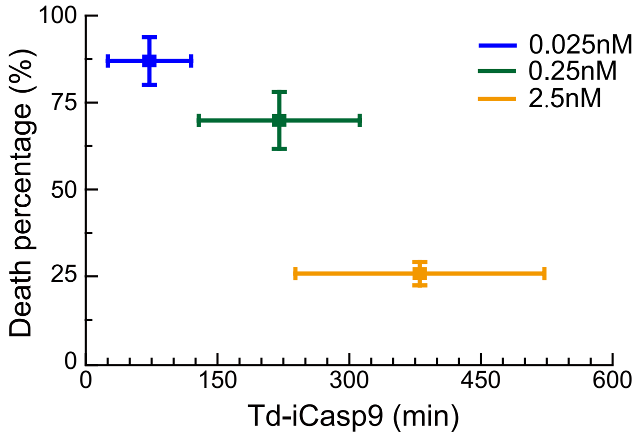


**Supplementary Fig. 5 Correlation between cell death and Td-iCasp9.** Death percentage is plotted against Td-iCasp9. Data are shown as mean ± standard deviation. Related to Figure 5

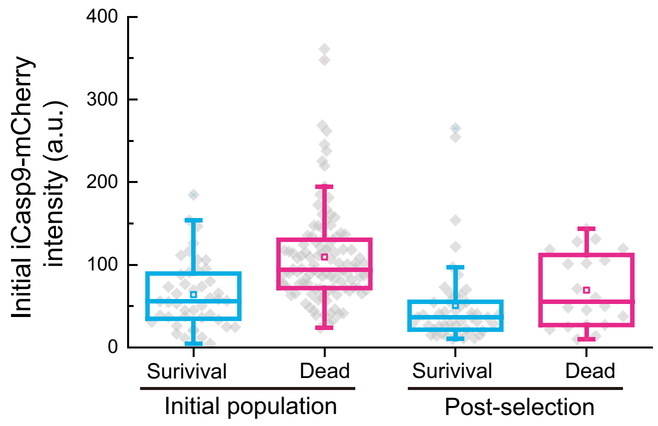


**Supplementary Fig. 6 Initial iCasp9-mCherry level in survival and dead cells.** Initial iCasp9-mCherry level in survival and dead cells after the first (initial population) and third round (post-selection) of inducer treatment. The boxes of the box plot represent the interquartile range between the first and third quartiles, whereas the whiskers represent the 95% and 5% values, and the squares represent the average. Related to Figure 6
